# Stimulus-dependent delay of perceptual filling-in by microsaccades

**DOI:** 10.1101/2024.10.16.618545

**Authors:** Max Levinson, Christopher C. Pack, Sylvain Baillet

## Abstract

Perception is a function of both stimulus features and active sensory sampling. The illusion of *perceptual filling*-in occurs when eye gaze is kept still: visual boundary perception may fail, causing adjacent visual features to remarkably merge into one uniform visual surface. Microsaccades–small, involuntary eye movements during gaze fixation–counteract perceptual filling-in, but the mechanisms underlying this process are not well understood. We investigated whether microsaccade efficacy for preventing filling-in depends on two boundary properties, color contrast and retinal eccentricity (distance from gaze center). Twenty-one human participants (male and female) fixated on a point until they experienced filling-in between two isoluminant colored surfaces. We found that increased color contrast independently extends the duration before filling-in but does not alter the impact of individual microsaccades. Conversely, lower eccentricity delayed filling-in only by increasing microsaccade efficacy. We propose that microsaccades facilitate stable boundary perception via a transient retinal motion signal that scales with eccentricity but is invariant to boundary contrast. These results shed light on how incessant eye movements integrate with ongoing stimulus processing to stabilize perceptual detail, with implications for visual rehabilitation and the optimization of visual presentations in virtual and augmented reality environments.

**Significance Statement:** To perceive, sense organs actively sample the environment—for example, by touching, sniffing, or moving the eyes. Visual sampling persists even when gaze is fixed on a single point: involuntary microsaccades continuously move the eye in small jumps. We investigated a previously documented observation that microsaccades prevent illusory fading of perceived visual boundaries during fixation. We discovered that despite being connected, microsaccades and fading are sensitive to different stimulus features. Boundaries separating surfaces with more distinct colors inherently took longer to fade. Boundaries closer to the center of vision also took longer to fade, but only because microsaccades were more effective. These findings reveal new insight into how pervasive sensory sampling delivers a stable and detailed perceptual experience.

## Introduction

When the eyes remain still for an extended period of time, visual boundaries seemingly fade from view. Boundary fading due to neuronal adaptation (Clarke & Belcher, 1962; Martinez-Conde & Macknik, 2017; Kohn, 2007) is the first stage of the illusory phenomenon of *perceptual filling-in*, whereby distinct visual regions appear to merge into a singular, uniform field (De Weerd et al., 1998; Weil & Rees, 2011). Despite persistent scientific interest in visual fading and filling-in as consequences of failed perceptual stability, a comprehensive understanding of their underlying mechanisms remains elusive.

An important clue comes from observations that microsaccades—small, rapid, and mostly involuntary eye movements that occur during periods of gaze fixation—inhibit filling-in (Clarke & Belcher, 1962; Martinez-Conde et al., 2006; Troncoso et al., 2008; McCamy et al., 2012; Costela et al., 2017). The role of microsaccades in everyday vision is a matter of ongoing debate (Kowler & Steinman, 1980; Martinez-Conde & Macknik, 2017; Poletti & Rucci, 2010), and it has been argued that this intriguing laboratory finding is an evolutionary artifact (Collewijn & Kowler, 2008; Rucci & Poletti, 2015). However, such controlled conditions can offer valuable insights into microsaccadic impacts on perception, especially related to the neural processes supporting perceptual stability. Understanding how microsaccades contribute to visual stability could inform the development of interventions for individuals with impaired visual function, such as those with age-related macular degeneration or other central vision loss conditions. Additionally, the insights gained from this study could help optimize visual displays in augmented and virtual reality environments in the context of naturalistic eye movement patterns.

The precise mechanism through which microsaccades counteract filling-in is not well understood. Crucially missing is an account of how stimulus characteristics—such as boundary eccentricity and contrast—interact with microsaccade efficacy for delaying filling-in, given that, for unclear reasons, stronger boundaries take longer to fade (Levinson & Baillet, 2022; Weil & Rees, 2011). It is often proposed that microsaccades serve to “refresh” visual stimuli (Engbert, 2006; Martinez-Conde et al., 2006), suggesting they generate cortical signals akin to visual stimulation but of reduced intensity. If so, microsaccades should be more effective over a more pronounced boundary. Alternatively, microsaccades might influence adapting neurons in unique ways, as they introduce minor retinal motion and alter the spatiotemporal configuration of the retinal image (Rucci & Victor, 2015). To examine the interplay between stimulus properties and microsaccade efficacy we developed a perceptual filling-in task based on the Uniformity Illusion (Otten et al., 2016) and analyzed microsaccade dynamics. Participants fixated their gaze until they perceived the merging of a central disk and the surrounding periphery. The circular boundary differed across trials along two axes of boundary strength: isoluminant color contrast and retinal eccentricity.

Using linear mixed modeling, we quantified how stimulus properties and microsaccades interact to influence perceptual stability. Our results demonstrate that microsaccade efficacy in delaying perceptual filling-in is influenced by boundary eccentricity but not by color contrast. Specifically, microsaccades were more effective at lower eccentricities, while color contrast independently extended the time required for perceptual filling-in. The findings suggest that microsaccades do not merely recreate initial stimulus features, but instead prevent perceptual filling-in by transiently refreshing visual signals in a manner dependent on cortical magnification. This study provides new insights into the mechanisms underlying perceptual stability and the role of incessant eye movements in visual processing, with implications for both basic neuroscience and clinical applications.

## Methods

### Participants

Twenty-four participants (14 females, age range: 19-45 years, all right-handed, including one author [ML]) took part in the study. All participants provided written informed consent before participation and were compensated $30 CAD per hour for their time. The study was conducted under the approval of the McGill University Health Centre Research Ethics Board (protocol 2021-7130). All participants were neurologically healthy and had normal or corrected-to-normal vision. Three participants were excluded before completing their first experimental session: two due to technical issues with the eye-tracking equipment and one due to difficulty maintaining fixation. Consequently, data from 21 participants were analyzed. One participant (subject 203) completed only 8 of the 9 task blocks.

The target sample size was determined by tripling the number of participants—8—included in the seminal study showing that microsaccades counteract visual fading (Martinez-Conde et al., 2006). Our final sample is larger than those in subsequent studies replicating this effect (Costela et al., 2017; McCamy et al., 2012; Troncoso et al., 2008).

### Stimuli and Task

The visual stimulus consisted of a central purple disk (the “center”) surrounded by a greyer rectangular background (the “periphery”). The entire display was contained within a 40.3 × 22.7-degree rectangular frame centered on the screen. The center and periphery were separated by a circular boundary, with a width equal to 12% of the center’s radius, allowing for a seamless transition between the two colors.

Participants completed 450 trials, divided into nine blocks of 50 trials each. At the start of each trial, participants pressed the spacebar to begin. A fixation target, designed to minimize fixational eye movements (Thaler et al., 2013), appeared for one second before the onset of the stimulus.

Participants were instructed to fixate on the fixation cross, minimize blinking, and press the spacebar once they perceived the center boundary fade and the two colored regions merge into one uniform color. Participants were informed that this was an illusion and that no physical changes would occur on the screen. The stimulus disappeared either 1 second after the spacebar press or automatically after 20 seconds from stimulus onset if no response was given. After each trial, dynamic colored noise was displayed for a duration equivalent to half of the stimulus presentation time to alleviate retinal fatigue and after-images.

Each trial block was defined by a specific combination of color contrast and boundary eccentricity, with block order randomized for each participant. This ensured that all trials within a block were identical, preventing interference from after-images of previous trials. Boundary eccentricity was determined by the radius of the center disk, measured in visual angle, with three sizes: 2 degrees (small), 4 degrees (medium), and 6 degrees (large). The stimulus color scheme was defined using an estimation of DKL color space (Derrington et al., 1984), enabling chromatic modulation measured in degrees around an isoluminant plane. In this DKL plane, all color values maintained a consistent radius of 0.07 and an elevation of 0 relative to a neutral grey baseline (RGB 77,77,77). Color contrast was quantified as the difference between the central and peripheral hues, specifically their DKL azimuth values. The center color was fixed at an azimuth of 270 degrees (RGB 77,75,83), while the peripheral hue varied across three azimuth values: 280 degrees (low contrast; RGB 83,73,82), 290 degrees (medium contrast; RGB 88,71,82), and 300 degrees (high contrast; RGB 92,69,82). RGB values were converted from DKL using the Computational Colour Science Toolbox 2e (Westland, 2021) and custom MATLAB code.

Each block included 50 trial: 40 “main” trials and 10 randomly distributed catch trials. Five of these catch trials were “replay” trials, which simulated the perceptual filling-in experience to validate participants’ accuracy in reporting filling-in events. During replay trials, the hues of the central and peripheral regions gradually merged into a uniform color over 2 seconds, with the onset of this effect occurring randomly between 4 and 6 seconds after stimulus presentation, following a uniform distribution.

The remaining five catch trials were “sharp” control trials, in which the boundary between the center and periphery was distinct, reduced to a single pixel’s width to create a sharp, well-defined edge. These trials aimed to confirm that a very pronounced boundary would be more resistant to fading, potentially leading to longer fixation times before participants perceived the merging of the two regions.

### Isoluminance Calibration

As a preliminary step, we ensured that the different stimulus colors were perceived as isoluminant by each participant. Isoluminance calibration was conducted using heterochromatic flicker photometry. During this procedure, a flickering circle with a radius of 200 pixels (approximately 4.2 degrees of visual angle) was displayed at the center of the screen. This circle alternated between the central hue (purple) and one of the peripheral hues at a frequency of 15 Hz, set against a grey background of fixed luminance (RGB 77,77,77). Participants adjusted the RGB values of the peripheral hue using the arrow keys on their keyboard, with goal of minimizing the flicker until it was either invisible or least noticeable.

This calibration was repeated for each of the three peripheral hues corresponding to the study’s three levels of color contrast. A majority of the 21 analyzed participants found the default peripheral colors to be isoluminant: 81% for low contrast, 76% for medium contrast, and 67% for high contrast. For participants who required adjustments, the changes were minimal, with average deviations from the default RGB values being low across all contrast levels: 1 ± 0 (low contrast), 1.4 ± 0.89 (medium contrast), and 1.43 ± 0.79 (high contrast).

This careful calibration ensured that any observed differences in filling-in time during the main task were due to the intended properties of the visual boundary and microsaccade dynamics, rather than variations in stimulus luminance.

### Experimental Procedure

The experiment consisted of three sessions, each scheduled on separate days within a two-week period. During the first session, participants received both written and oral instructions, followed by a practice session of 10 trials at medium contrast and eccentricity to familiarize them with the task.

Participants completed three task blocks per session. They sat in a dimly lit room with their head stabilized in a chinrest, positioned 60 cm away from a 27-inch monitor (resolution: 2560 × 1440 pixels; refresh rate: 60 Hz). Visual stimuli were generated using *Psychtoolbox3* (Brainard, 1997) in MATLAB R2020b (The MathWorks Inc., 2020). Eye movements were recorded at a 1000-Hz sampling rate with an EyeLink 1000 Plus infrared video eye tracker (SR Research) positioned in front of the monitor. Each block began with an eye-tracking calibration and validation process. The first 9 participants underwent a 5-point calibration, while the subsequent 12 participants received a more extensive 9-point calibration. No systematic differences were observed between the two calibration groups.

Participants initiated each trial by pressing the spacebar, allowing a self-paced approach to maximize comfort and readiness. Eye gaze was continuously monitored, and trials were automatically restarted if gaze deviation exceeded 2 degrees from the fixation point for more than 50 ms (3 continuous frames).

### Eye Movement Analysis

Trials were excluded if they were shorter than 2 seconds (filling-in time [FT] < 2 seconds) to prevent accidental early responses, if median gaze displacement exceeded 1 degree from the fixation cross, or if a blink or a large saccade (velocity > 30 degrees/second velocity, detected automatically by the EyeLink control computer) occurred within 300 ms before the button press. The 300-ms cutoff was crucial to exclude ocular events potentially related to motor execution rather than visual perception. Of 7560 main trials, 6712 met these criteria and were analyzed further.

Microsaccades were detected using a refined velocity-based algorithm adapted from Engbert & Kliegl (2003). Eye gaze data from each trial were converted into visual angle velocity using a 31-ms sliding window. To minimize artifacts, data within 150 ms before and after each blink were excluded. Microsaccades were identified as sequences where, for at least eight consecutive timepoints, velocity exceeded a threshold based on the standard deviation above noise (Engbert & Kliegl, 2003) in both eyes (binocular detection). Microsaccades separated by less than 12 ms were merged into a single event. Additional criteria required that eye movement direction did not vary by more than 15 degrees per millisecond and that the total amplitude fell between 3 arcmin and 2 degrees. Eye movements smaller than 3 arcmin were classified as ocular drift or noise, while movements larger than 2 degrees were classified as large saccades.

To determine the optimal noise threshold for each participant, we manually reviewed detected microsaccades in every trial. The thresholds for noise multipliers were calibrated based on trial-by-trial manual review to ensure accuracy in detecting true microsaccades versus noise. Eighteen participants were assigned a noise multiplier of 5, while three required a multiplier of 6.

To qualitatively examine microsaccade dynamics in relation to perceptual filling-in, we assembled a vector of microsaccade onsets for each trial, aligning the spacebar press (indicating filling-in) to time zero. These vectors were summed across trials to calculate microsaccade frequency as a rate per second, using a causal windowing function (Engbert, 2006) with a window length of 1001 ms and an alpha value of 1/100—similar to methods used in neuronal firing rate analysis (Dayan & Abbott, 2001). Trials with blinks or large saccade within 2 seconds of the button press were excluded (543 excluded trials). For visualization, we normalized microsaccade rates so that each participant’s average rate during the baseline period (−5 to -3 seconds before button press) matched the study population’s average baseline rate.

Ocular drift was measured using the “retinal slip” methodology (Engbert & Mergenthaler, 2006), which calculates the distance of gaze traversal in visual angle over time. The screen was divided into a grid of 0.01 × 0.01-degree squares, approximately representing the diameter of a cone’s receptive field. We tracked the number of squares traversed in each 50-ms segment of a trial, converting the count into a rate of degrees per second. Timepoints during or immediately (10 ms) before or after a microsaccade were excluded to isolate ocular drift from microsaccadic movements.

### Linear Mixed Modeling of Filling-in Time

We used linear mixed models to predict the time taken for perceptual filling-in in each trial, considering various stimulus attributes and eye movements. The models were built using the *lme4 v11*.*1*.*35*.*1* package (Bates et al., 2015) in *R Statistical Software v4*.*3*.*2* (R Core Team, 2023). Our approach involved systematic model construction and parameter selection, testing specific hypotheses without overcomplicating the models.

Filling-in times (FT) were log-transformed to improve model fitting. Continuous predictor variables were standardized by zero-centering and scaling by their standard deviations. We used restricted maximum likelihood estimation (REML) with the *BOBYQA* optimization algorithm for model fitting. Parameters were considered statistically significant if their Bonferroni-corrected confidence intervals (excluding the intercept) did not include 0 (see *Statistical Analysis*). Post-hoc analyses were performed to determine if removing them improved model fit, as indicated by a lower Bayesian information criterion (BIC). Parameters whose removal did not improve model fit were retained for inclusion in future models.

Model A (Table S1) was designed to predict FT based solely on stimulus properties, without considering eye movements. Included trial-by-trial variables were stimulus contrast, eccentricity, and trial number, which helped assess any changes in FT across the block. The *lme4* syntax was defined as:

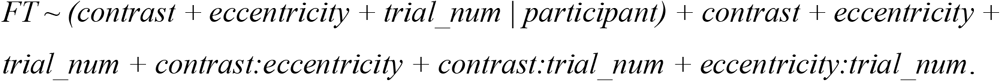

Model B (Table S2) incorporated the significant parameters from Model A, adding main effects of four eye movement variables: microsaccade presence (whether microsaccades occurred during a trial), number of blinks, and average ocular drift. We focused on microsaccade presence (rather than count or rate) for two reasons: 1) to determine if microsaccades prolonged FT in general, and how this effect interacted with contrast or eccentricity, and 2) to avoid potential spurious correlations between longer trial durations and increased spontaneous eye movements, which are challenging to interpret. The *lme4* syntax for Model B was:

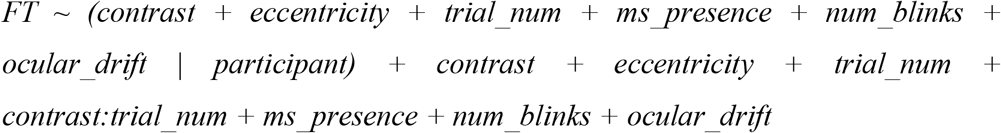

The primary model (Table 1) included fixed and random main effect variables identified from Model B, plus their fixed-effect interactions. The critical tests of interest were the interaction effects between microsaccade presence and stimulus contrast or eccentricity. The *lme4* syntax was:

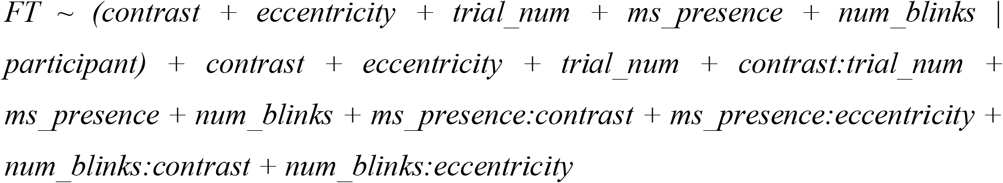

Two additional models (Table 2) were used to further verify whether the effects of contrast and eccentricity on FT depended on microsaccade occurrence. We separately analyzed trials without any microsaccades or blinks and trials with at least one microsaccade but no blinks. To balance trial counts across both models for each participant, we randomly excluded trials from the model that contained more trials, resulting in 630 total trials for each model (9.4% of the total). Both models used the following *lme4* syntax:

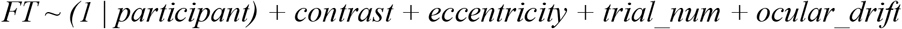

Given the reduced trial counts, random slopes were not fitted, and we used uncorrected 95% confidence intervals to minimize the risk of false negatives.

**Table 1.**
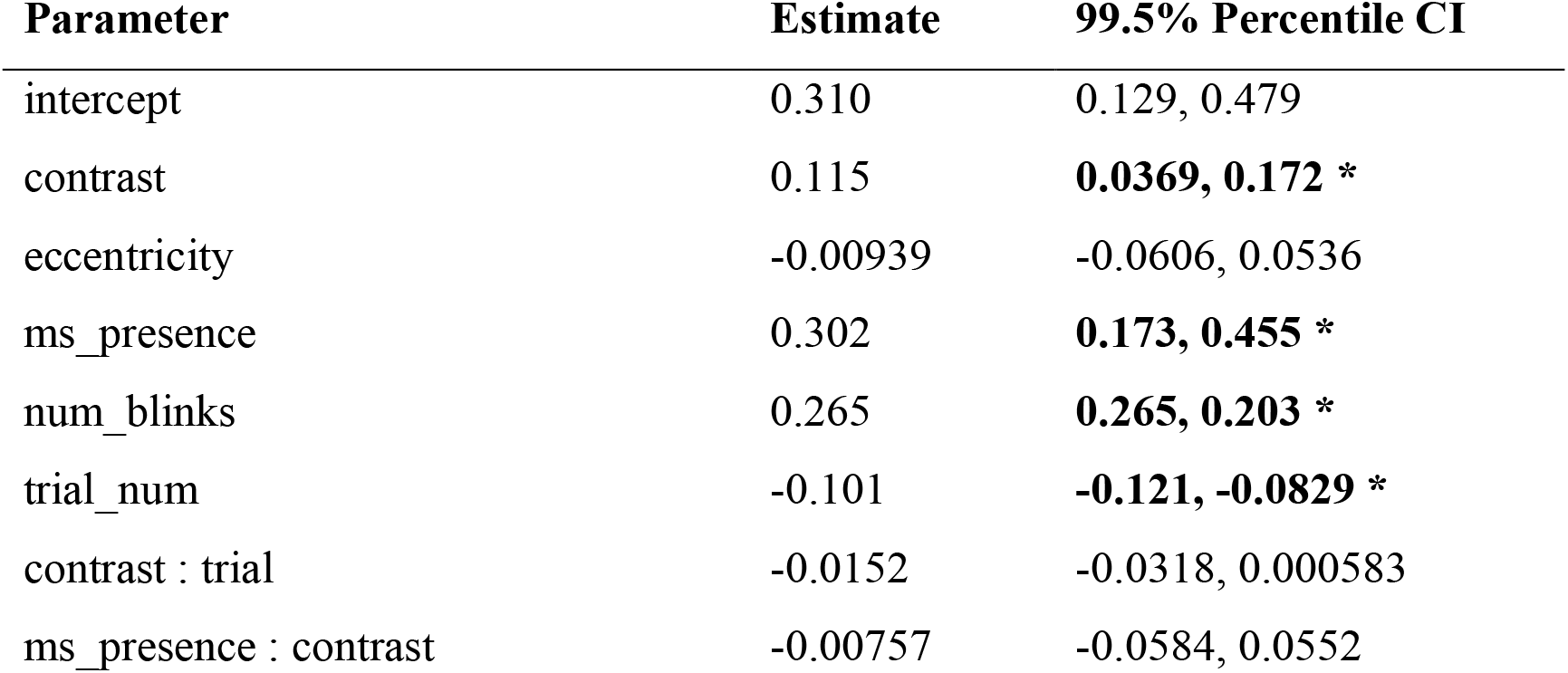
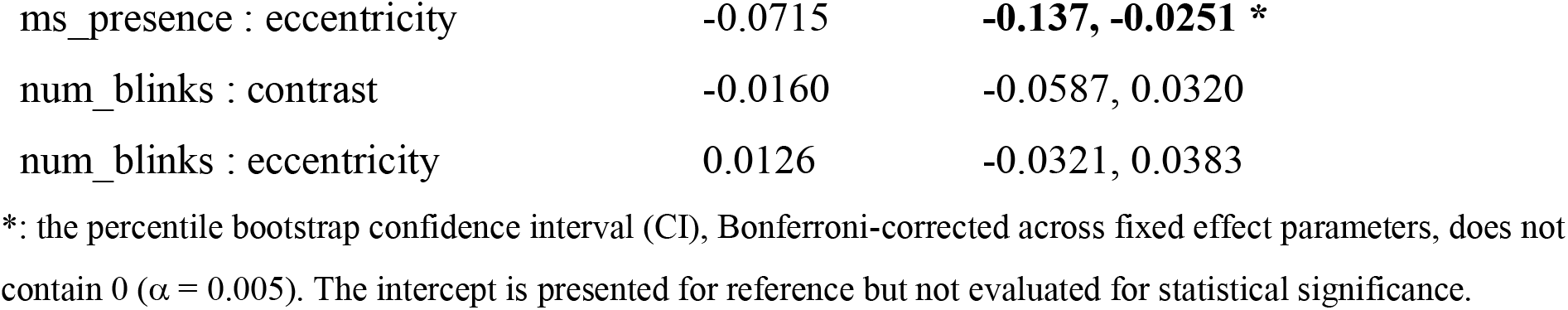
Fixed effects from a linear mixed-effect model of log-transformed filling-in time.

| Parameter | Estimate | 99.5% Percentile CI |
| --- | --- | --- |
| intercept | 0.310 | 0.129, 0.479 |
| contrast | 0.115 | **0.0369, 0.172 \*** |
| eccentricity | -0.00939 | -0.0606, 0.0536 |
| ms_presence | 0.302 | **0.173, 0.455 \*** |
| num_blinks | 0.265 | **0.265, 0.203 \*** |
| trial_num | -0.101 | **-0.121, -0.0829 \*** |
| contrast : trial | -0.0152 | -0.0318, 0.000583 |
| ms_presence : contrast | -0.00757 | -0.0584, 0.0552 |
| ms_presence : eccentricity | -0.0715 | **-0.137, -0.0251 \*** |
| num_blinks : contrast | -0.0160 | -0.0587, 0.0320 |
| num_blinks : eccentricity | 0.0126 | -0.0321, 0.0383 |

\*: the percentile bootstrap confidence interval (CI), Bonferroni-corrected across fixed effect parameters, does not contain 0 ( $\alpha = 0.005$ ). The intercept is presented for reference but not evaluated for statistical significance.

**Table 2.** Fixed effects from a comparative linear mixed-effect model analysis of microsaccade-dependent filling-in time.

| Parameter | microsaccade trials |  | no-microsaccade trials |  |
| --- | --- | --- | --- | --- |
|  | Estimate | 95% Percentile CI | Estimate | 95% Percentile CI |
| intercept | 0.498 | 0.363, 0.620 | 0.176 | 0.0636, 0.279 |
| contrast | 0.156 | **0.0941, 0.202 \*** | 0.166 | **0.0851, 0.225 \*** |
| eccentricity | -0.0876 | **-0.131, -0.0350 \*** | -0.00389 | -0.0305, 0.0310 |
| trial_num | -0.101 | **-0.130, -0.0722 \*** | -0.0762 | **-0.100, -0.0552 \*** |
| ocular_drift | 0.104 | **0.000418, 0.170 \*** | 0.0453 | **0.0183, 0.129 \*** |

Linear mixed models evaluated filling-in time (FT) in two distinct subsets of trials: those with at least one microsaccade occurring (630 trials, left columns) and those devoid of microsaccades (630 trials, right columns). \*: the percentile bootstrap confidence interval (CI) does not contain 0 ( $\alpha = 0.05$ ). The intercept is presented for reference but not evaluated for statistical significance.

### Linear Mixed Modeling of Microsaccade Generation

We evaluated the influence of boundary contrast, eccentricity, and trial number on microsaccade generation, measured as the trial-wise microsaccade rate. The model was formulated in *lme4* syntax as:

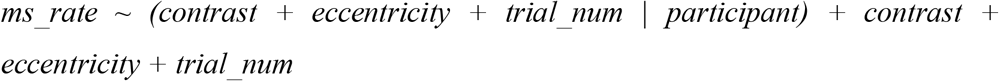

### Linear Mixed Modeling of Immobilization Time

We also conducted linear mixed modeling to analyze immobilization time, defined as the duration between the last microsaccade in a trial and the button press indicating perceptual filling-in. To ensure relevance to the perceptual experience, we included only trials where the last microsaccade occurred more than 300 ms before the button press (4692 trials). This criterion was crucial to exclude eye movements that might occur after the perceptual event but before the motor response, which could potentially confound our analysis (Betta & Turatto, 2006). The model for immobilization time was structured as:

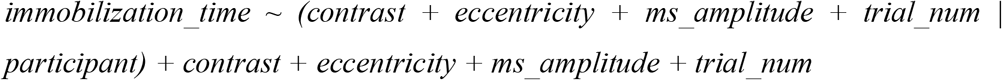

### Statistical Analysis

Average filling-in times for main trials and “sharp” catch trials were compared using a Wilcoxon signed-rank test. To determine the statistical significance of fixed effect parameters in all linear mixed models, we used case-bootstrapping with the *lmeresampler v0*.*2*.*4 R* package (Loy et al., 2023). This approach was chosen because it validates parameter estimates with minimal assumptions regarding data distribution (Leeden et al., 2008). By resampling participants’ data with replacement, we could derive more robust confidence intervals for each fixed effect parameter, thus improving the reliability of our inferences about microsaccadic effects. For each model, we performed 10,000 resampling iterations, resampling individual participants’ data with replacement, to derive percentile confidence intervals for each fixed effect parameter.

## Results

### Both Eccentricity and Color Contrast Prolong Filling-in Time

Twenty-one human participants completed a perceptual task to assess isoluminant color filling-in across the boundary between a central disk and its surrounding periphery (Fig. 1a). Filling-in was reported in the majority of main trials (95.4% ± 4.75; mean ± standard deviation across participants), with an average filling-in time (FT) of 7.51 ± 0.48 seconds (mean ± standard error).

**Fig. 1.**
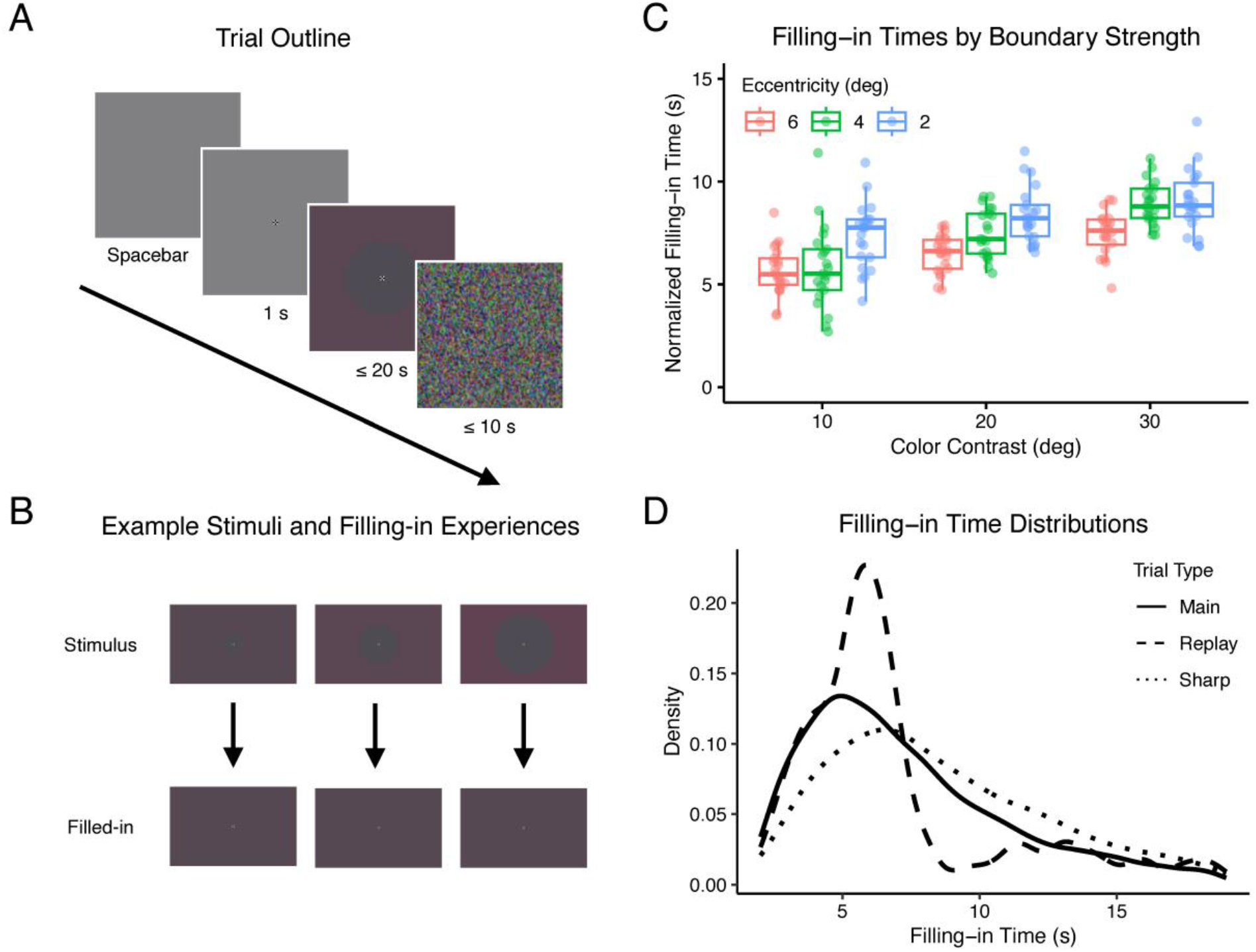
Experiment Design and Behavior. **(A)Sequence of events in a typical trial**. Participants initiated each trial by pressing the spacebar, and the visual stimulus persisted until a second spacebar press or a maximum of 20 seconds elapsed. Dynamic noise followed each trial to mitigate retinal afterimages. **(B)Stimulus transition and perceptual effect**. The stimulus included a central disk varying in radius and with three possible color contrasts (top row). After boundary fading, the central and peripheral regions perceptually merged, creating a uniform color field (bottom row). **(C)Normalized filling-in time across main trials**. plotted against the boundary’s color contrast and eccentricity. Each data point represents an individual participant’s experience. Boxplots show median, 25^th^ and 75^th^ percentiles across individual subject means. **(D)Distribution of filling-in times across different trial types**. Control trials with sharp boundaries (dotted line) typically resulted in prolonged FT compared to main trials (solid line), while replay trials (dashed line) showed clustered FT around the simulated effect (6-8 seconds).

The results showed a clear relationship between boundary strength and filling-in time (FT). Both higher color contrast and lower eccentricity boundaries increased FT (Fig. 1b, 1c; Table S1; linear mixed modeling, *α* = 0.0083). No significant interaction was found between color contrast and eccentricity in their effects on FT. A trend was also observed where FT decreased as the task block progressed, and the effect of color contrast on FT diminished over time (Table S1).

In the 45 controlled “replay” catch trials, FT was concentrated around the time when the stimulus became visually uniform (6-8 seconds; Fig. 1d), with reduced variability across stimulus conditions (Fig. S1a). In the set of 45 “sharp” trials intended to extend filling-in time, FT exceeded that of main trials for all participants (9.12 ± 0.55 seconds; Wilcoxon signed-rank test, *W* = 0, *p* < 0.001, Fig. 1d), across all stimulus conditions (Fig. S1b). The consistent results across replay and sharp trials indicate that participants’ responses accurately reflected their perceptual experience of filling-in.

### Eccentricity Only Predicts Filling-in time When Microsaccades Occur

The primary analysis confirmed two key assumptions regarding microsaccade behavior: 1) detected microsaccades conformed to the classic “main sequence” relationship, which describes the correlation between magnitude and peak velocity (Bahill et al., 1975; Fig. 2a), and 2) microsaccade rate exhibited a notable dip approximately 600 ms before participants reported perceptual filling-in (Costela et al., 2017; Martinez-Conde et al., 2006; McCamy et al., 2012; Troncoso et al., 2008; Fig. 2b). Microsaccade rate profiles remained consistent across all nine stimulus conditions (three levels of color contrast and three levels of eccentricity), with no significant deviations beyond random variation (detailed rate curves in Fig. S2).

**Fig. 2.**
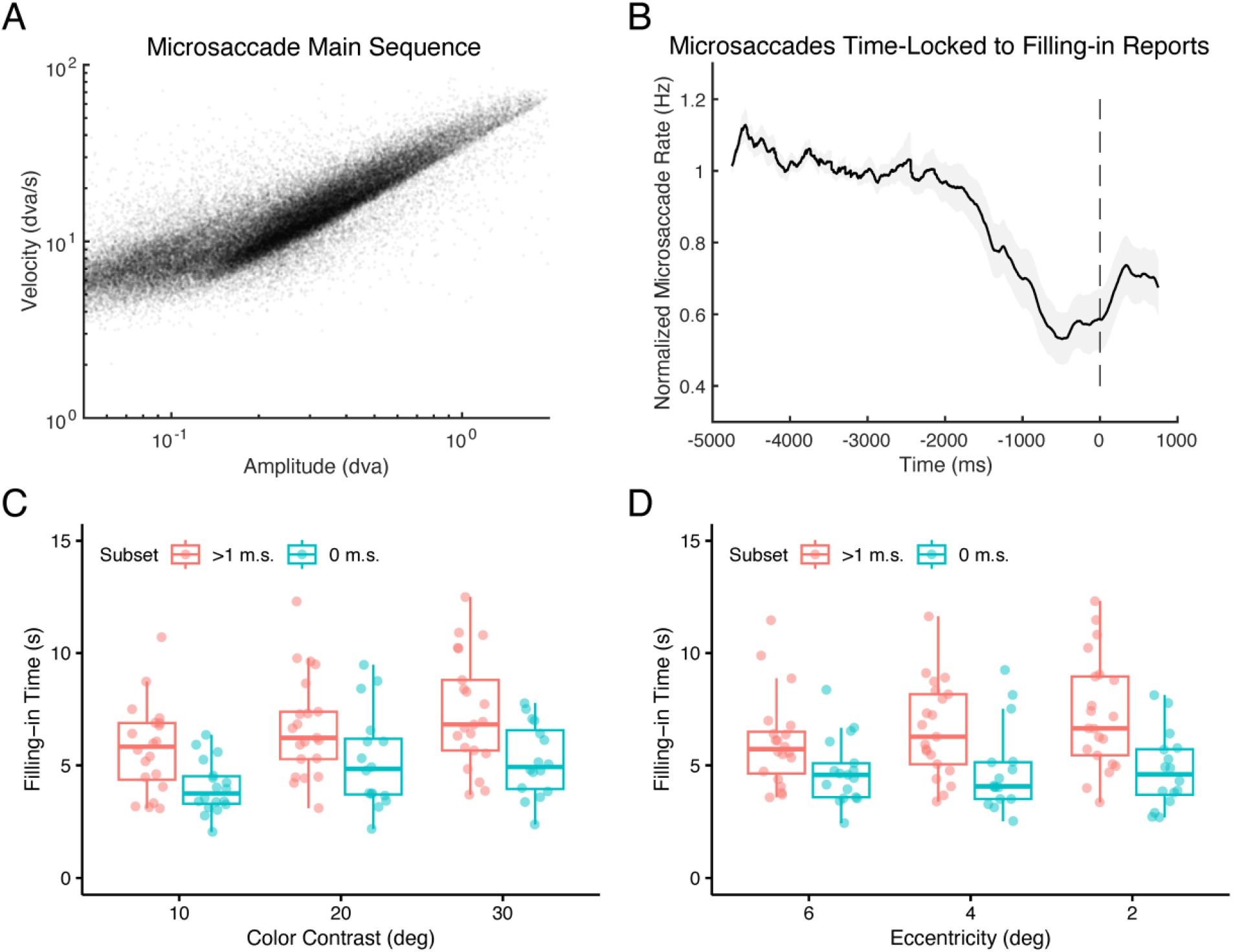
Microsaccade Dynamics and Their Influence on Filling-in Time. **(A)**Relationship between amplitude and peak velocity of individual microsaccades during main trials, demonstrating adherence to the classic main sequence relationship. **(B)**Average normalized microsaccade rate in the period leading up to participants’ report of perceptual filling-in. **(C)**Filling-in time as a function of color contrast, comparing trials with at least one microsaccade (red) to trials without microsaccades (blue). **(D)**Filling-in time as a function of boundary eccentricity, comparing trials with at least one microsaccade (red) to trials without microsaccades (blue). Each data point represents an individual participant. Boxplots show the median, 25^th^, and 75^th^ percentiles of participants’ means.

The comprehensive linear mixed model outcomes, with bootstrap-derived confidence intervals (*α*= 0.005), are detailed in Table 1 (see Tables S1 and S2 for precursor models used for parameter selection). Our findings demonstrate that filling-in time was significantly prolonged in trials where microsaccades occurred, as well as in trials with increased blinks. Among the interactions between stimulus properties and ocular movements, the interaction between microsaccade presence and boundary eccentricity was the only significant effect. Specifically, the impact of boundary eccentricity on FT was significantly amplified in the presence of microsaccades, while color independently influenced FT. These findings remained consistent when recalibrating eccentricity values using a logarithmic conversion based on cortical magnification. This conversion translates visual field eccentricity (*E*) into cortical distance (*d*, in mm) from the retinotopic representation corresponding to 10 degrees of visual angle (Engel et al., 1997): d = log(E) / 0.063 - 36.54.

We further examined whether boundary strength, manipulated through color contrast and eccentricity, influenced perceptual filling-in time independently of microsaccades. Trials were divided based on ocular stability: those with no microsaccades or blinks, and those with at least one microsaccade but no blinks. As hypothesized, both color contrast and eccentricity significantly predicted FT in trials where microsaccades occurred (Table 2, left column; Fig. 2c-d; *α* = 0.05). Interestingly, in trials without microsaccades—indicating stable fixation—eccentricity had no effect on FT (Table 2, right column; Fig. 2d). However, color contrast continued to predict FT even under stable fixation (Fig. 2c).

To test an alternative hypothesis—that stronger visual boundaries prolong FT by increasing the rate of microsaccade generation rather than their efficacy—we analyzed the relationship between microsaccade rate and boundary properties. Contrary to this hypothesis, we found no significant association between microsaccades rate and either color contrast or eccentricity (Table 3; Fig. 3a; *α* = 0.0167).

**Table 3.** Fixed effects from a linear mixed-effect model of log-transformed microsaccade rate.

| Parameter | Estimate | 98.33% Percentile CI |
| --- | --- | --- |
| intercept | -0.0845 | -0.307, 0.128 |
| contrast | -0.000870 | -0.0449, 0.0426 |
| eccentricity | 0.0350 | -0.00397, 0.0723 |
| trial_num | 0.134 | **0.0893, 0.175 \*** |

\*: the percentile bootstrap confidence interval (CI), Bonferroni-corrected across fixed effect parameters, does not contain 0 ( $\alpha = 0.0167$ ). The intercept is presented for reference but not evaluated for statistical significance.

**Fig. 3.**
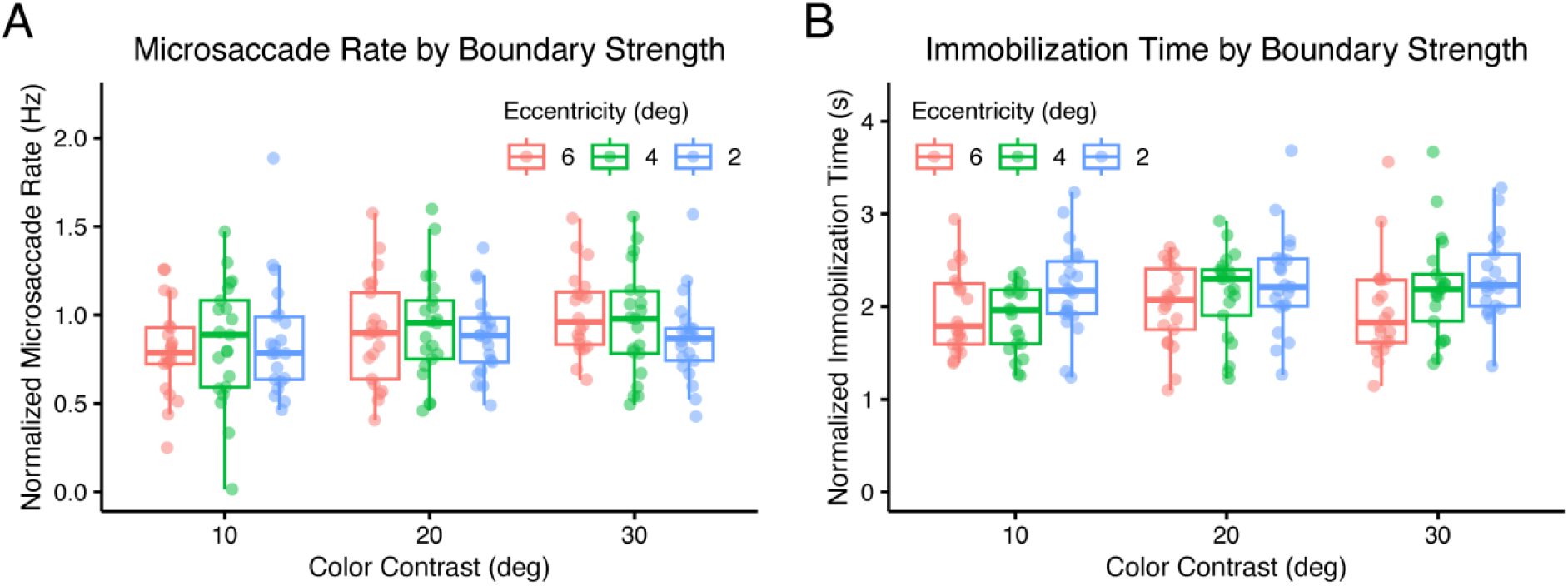
Microsaccade Rates and Immobilization Times in Relation to Visual Stimulus Parameters. **(A)**Normalized microsaccade rate throughout trials as a function of color contrast and boundary eccentricity. **(B)**Normalized immobilization time as a function of color contrast and boundary eccentricity. Immobilization time represents the duration of stable fixation after the last microsaccade until the report of perceptual filling-in. Each data point represents an individual participant. Boxplots show the median, 25^th^, and 75^th^ percentiles across participants’ means.

### Eccentricity, but Not Color Contrast, Influences Microsaccadic Efficacy

Another way to evaluate microsaccade efficacy is to quantify the additional fixation time required for perceptual filling-in following each microsaccade. According to slow adaptation hypotheses of boundary fading, neural signals gradually diminish until the visual boundary fades from perception. Microsaccades counteract this adaptation by restoring part of the diminished signal, delaying fading and subsequent filling-in. The amount of signal restoration—and hence the delay—would be expected to increase with more effective microsaccades, such as those occurring at smaller boundary eccentricities. Although we cannot determine the exact microsaccade-induced delay or signal increment, we can infer how reduced eccentricity extends this delay by analyzing “immobilization time”--the duration from the last microsaccade to the report of filling-in, reflecting the time the eyes must remain stationary for filling-in to occur.

As hypothesized, immobilization time was significantly associated with eccentricity but not with color contrast, as revealed by a linear mixed model (Table 4; Fig. 3b; *α* = 0.0167). Average immobilization times were 2220 ± 242 ms for the 2-degree eccentricity condition, 2057 ± 229 ms for 4 degrees, and 1903 ± 184 ms for 6 degrees. Interestingly, the amplitude of the last microsaccade did not predict immobilization time, suggesting that the extent of retinal image displacement does not directly affect microsaccade efficacy in delaying filling-in. Immobilization times also tended to decrease as trials progressed within a block. It is important to note that the reported immobilization times include not only the delay induced by microsaccades but also any residual adaptation period and motor response time.

**Table 4.**
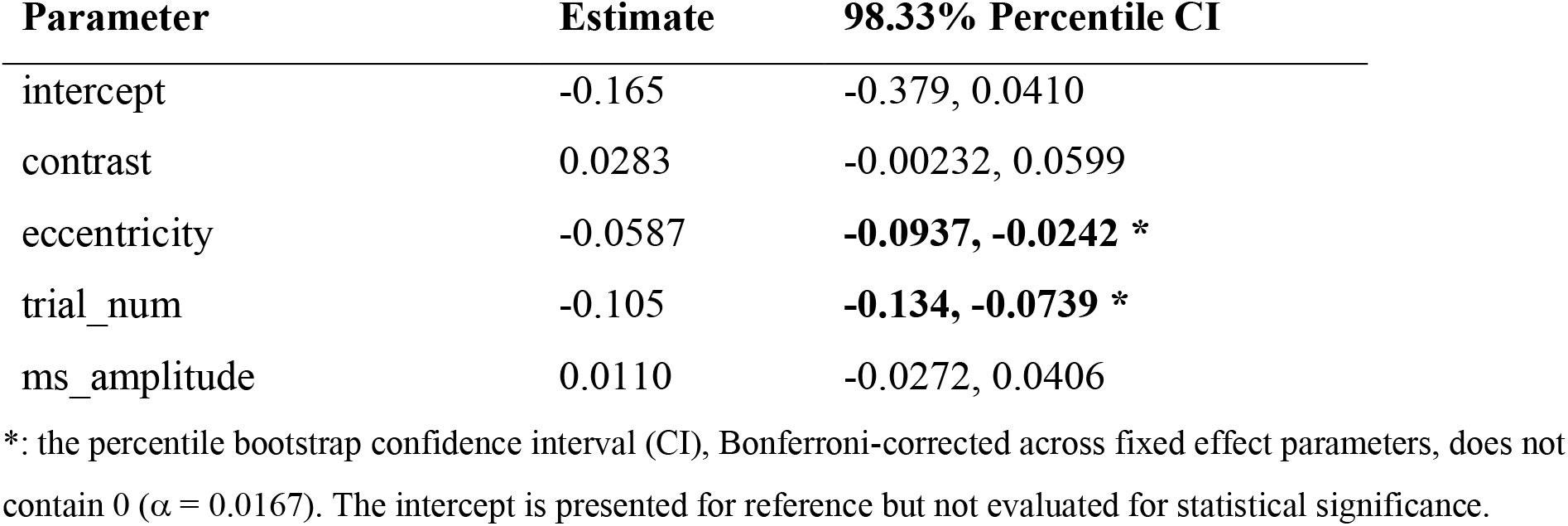
Fixed effects from a linear mixed-effect model of log-transformed immobilization time.

| Parameter | Estimate | 98.33% Percentile CI |
| --- | --- | --- |
| intercept | -0.165 | -0.379, 0.0410 |
| contrast | 0.0283 | -0.00232, 0.0599 |
| eccentricity | -0.0587 | **-0.0937, -0.0242 \*** |
| trial_num | -0.105 | **-0.134, -0.0739 \*** |
| ms_amplitude | 0.0110 | -0.0272, 0.0406 |

\*: the percentile bootstrap confidence interval (CI), Bonferroni-corrected across fixed effect parameters, does not contain 0 ( $\alpha = 0.0167$ ). The intercept is presented for reference but not evaluated for statistical significance.

## Discussion

During voluntary gaze fixation, microsaccades play a crucial role in maintaining visual perception by preventing the fading of visual boundaries that would otherwise lead to perceptual filling-in. Despite their significance, the mechanisms underlying how microsaccades modulate this process remain poorly understood. This study demonstrates that the efficacy of microsaccades in delaying perceptual filling-in is influenced by retinal eccentricity but is independent of color contrast, offering new insights into the mechanisms of perceptual stability. Specifically, we found that higher color contrast independently increased the time required for perceptual filling-in (filling-in time, FT), while microsaccades mitigated boundary fading equally across varying contrast levels. Conversely, lower boundary eccentricity increased the efficacy of microsaccades, whereas FT remained consistent across different eccentricities when no microsaccades were present. These conclusions were supported by a series of linear mixed model analyses that revealed: 1) a significant interaction between microsaccade occurrence and boundary eccentricity, but not color contrast, in predicting FT; 2) a significant effect of eccentricity on FT when microsaccades were present, but not in trials without microsaccades; and 3) a significant impact of eccentricity, but not color contrast, on immobilization time—the stable fixation period between the last microsaccade and the report of filling-in.

The lack of sensitivity of FT to eccentricity in the absence of microsaccades, despite the established relationships between eccentricity, visual acuity, and cortical magnification (Duncan & Boynton, 2003), suggests that visual adaptation of boundaries may occur uniformly across the visual field.

This finding aligns with previous studies (Bachy & Zaidi, 2014a; Greenlee et al., 1991) but highlights an indirect role of eccentricity in modulating the impact of microsaccades on adaptation.

Two secondary findings emerged regarding the progression of trials within each block. First, we observed a decrease in both filling-in time and immobilization time as trials progressed, which we attribute to practice effects resulting in faster motor responses. Second, we found that the effect of trial number on FT exhibited a negative interaction with color contrast, suggesting that the influence of contrast on FT diminished over time. Although this interaction did not meet the strict statistical threshold, excluding it from the model led to poorer performance, suggesting its potential relevance. One plausible explanation is that visual neurons gradually adapt to the higher-level, sustained visual context of a trial block, normalizing their responses over time (Webster, 2011).

Our study provides significant new insight into the mechanisms underlying microsaccades’ role in perceptual filling-in. We demonstrated that microsaccades do not simply resample the initial visual stimulus, as their efficacy in preventing filling-in varies with boundary eccentricity but is unaffected by color contrast. This finding suggests that microsaccades serve a more complex role than merely refreshing the visual input. The relationship between microsaccade efficacy and eccentricity can likely be attributed to cortical magnification and receptive field size—stimuli at higher eccentricities are processed by fewer cortical neurons with larger receptive field sizes (Daniel & Whitteridge, 1961). Consequently, at higher eccentricities fewer neurons are activated by the subtle retinal shifts induced by microsaccades (Donner & Hemilä, 2007). This hypothesis has been proposed previously (Clarke & Belcher, 1962) and could also be related to variations in retinal ganglion cell responses across eccentricities (Bachy & Zaidi, 2014b). These findings imply that certain visual field asymmetries in behavior may be entirely due to fixational eye movement dynamics. This could be of clinical relevance for optimizing visual rehabilitation strategies for individuals with central vision loss or age-related macular degeneration, where enhancing the stability of central visual representations is critical.

Interestingly, we found no correlation between microsaccade amplitude and efficacy. While larger amplitude microsaccades would theoretically stimulate more neurons, we found that this broader stimulation does not necessarily enhance filling-in prevention, consistent with prior findings (McCamy et al., 2014). It is possible that the critical neural population consists of cells with receptive fields at the boundary before or after a microsaccade, rather than all cells activated during the eye movement. Alternatively, the concept of corollary discharge, in which microsaccade generation modulates gain in visual cortex before the movement occurs (Chen et al., 2015), could offer a mechanism for the refreshing effect of microsaccades. This boost may be more pronounced at smaller eccentricities due to cortical magnification and may not vary with contrast. If this is the case, the actual retinal image shift may be irrelevant to boundary fading—an intriguing hypothesis that warrants further investigation.

We also found that microsaccade efficacy was insensitive to isoluminant color contrast, reinforcing the idea that microsaccades do not simply replicate the initial visual stimulus. This suggests that the feature defining the visual boundary and its adaptation dynamics does not modulate how microsaccades prevent such adaptation. Future research should investigate whether this invariance to color contrast extends to other visual properties, such as luminance. Such work would be essential for delineating the neural processes and brain regions involved in adaptation and microsaccadic counteraction. Given that neurons in the primary visual cortex (V1) process contours formed from both luminance and isoluminant color edges similarly (De & Horwitz, 2022; Hamburger et al., 2007), it seems unlikely that microsacadic efficacy would vary based on these properties. Instead, the mechanism may originate in motion-processing pathways, which are known to be invariant to luminance and color contrast for rapid retinal motion (Hawken et al., 1994), or through corollary discharge as discussed.

Our experimental design has certain limitations. Trials within each block were not randomized by contrast and eccentricity, potentially introducing non-stimulus-related effects, such as arousal fluctuations. However, the consistent relationship between FT and both contrast and eccentricity across participants makes this unlikely. Additionally, although replay control trials verified task accuracy, they did not fully mimic the gradual perceptual filling-in experience. Sudden changes in physical contrast can abruptly remove stimuli from conscious perception (May et al., 2003) due to normalization of neuronal firing rates to the current visual input. A reduction of neural firing below baseline can even produce opposite afterimages (Bachy & Zaidi, 2014a, 2014b), which would further differentiate the replay and filling-in experiences. Indeed, participants often reported the replay stimulus as inverting—first appearing uniform and then revealing the center disk in the opposite color. Nevertheless, our intention for including a replay condition was only to verify task engagement.

Lastly, the narrow range of color contrasts sampled may have limited our ability to detect subtle relationships between contrast and microsaccades. Although our findings suggest that any potential contrast effect on microsaccadic adaptation prevention is minimal compared to the effect on adaptation itself, further research using a broader contrast range is needed. Moreover, the industry-standard eye tracker used may not detect the smallest microsaccades (<15 arcmin; Collewijn & Kowler, 2008). Although trials without detected microsaccades were labeled accordingly, it is possible that undetected eye movements were present. Even so, our robust findings suggest that such small movements would be ineffective at preventing filling-in, raising the question of whether a minimum microsaccade magnitude is required to influence boundary fading.

## Conclusions

The present findings shed light on how fixational eye movements modulate the initial stage of perceptual filling-in—boundary fading. While higher color contrast inherently increases filling-in time, microsaccades specifically mediate the similar influence of boundary eccentricity. This suggests that microsaccades prevent boundary fading via transient boundary stimulation that scales with cortical magnification but is invariant to boundary contrast. Together, these findings offer a foundation for future research aimed at identifying the precise neurophysiological mechanisms underlying the interaction between eye movements and visual perception, with potential applications in both clinical and technological domains.

## Acknowledgments

This research was supported by the following sources of funding:

Quebec Bio-imaging Network project 12.49 (ML)

National Sciences and Engineering Research Council of Canada discovery grant RGPIN-2020-06889 (SB)

Canadian Institutes of Health Research Canada Research Chair Tier 1; CRC-2017-00311 (SB) National Institutes of Health grant R01-EB026299 (SB)

## Author contributions

Conceptualization: ML, CCP, SB

Methodology: ML, CCP, SB

Software: ML

Resources: CCP

Investigation: ML

Formal analysis: ML

Data curation: ML

Visualization: ML

Project administration:

ML Supervision: SB

Writing—original draft: ML

Writing—review & editing: ML, CCP, SB

Funding acquisition: ML, SB

## Data and materials availability

All data needed to evaluate the conclusions in the paper are available in the main text or the supplementary materials. All experimental and analysis code, raw behavioral and eyetracking data, and processed data files are available in a public repository: https://doi.org/10.17605/OSF.IO/KVT5A.

## List of Supplementary Materials

Figs. S1 to S2

Tables S1 to S2

## Notes

**Conflict of Interest:** The authors declare no competing financial interests.

https://doi.org/10.17605/OSF.IO/KVT5A

### Competing Interest Statement

The authors have declared no competing interest.

