## Supplementary Materials for "Stimulus-dependent delay of perceptual filling-in by microsaccades"

Max Levinson *et al.*

Supplementary Materials

Figs. S1 to S2
Tables S1 to S2

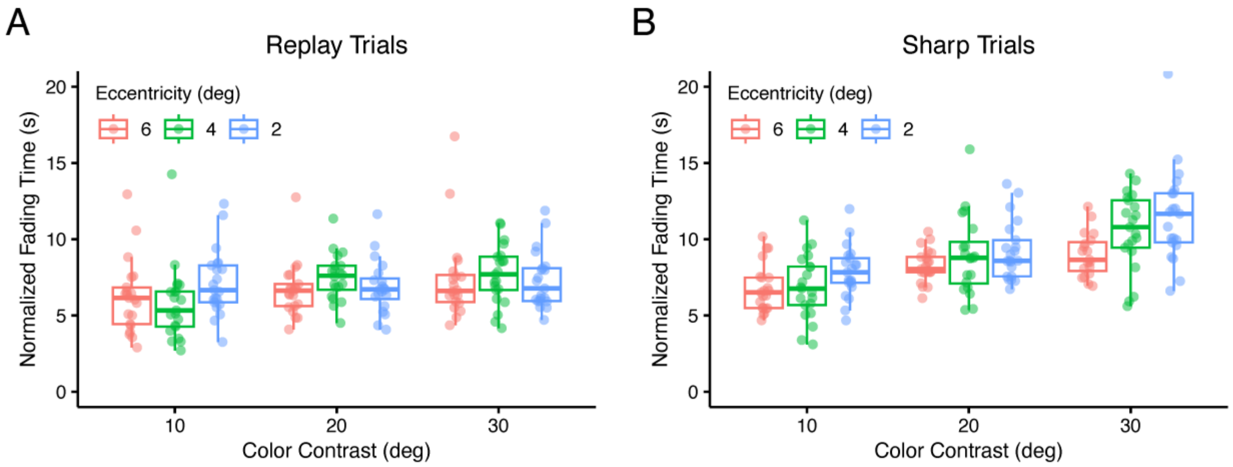


**Fig. S1. Catch trial filling-in times.** Normalized filling-in times for two categories of catch stimuli: **(A)** replay of the filling-in experience and **(B)** sharp, 1-pixel width boundaries between center and periphery. Boxplots show median, 25$^{th}$ and 75$^{th}$ percentiles.

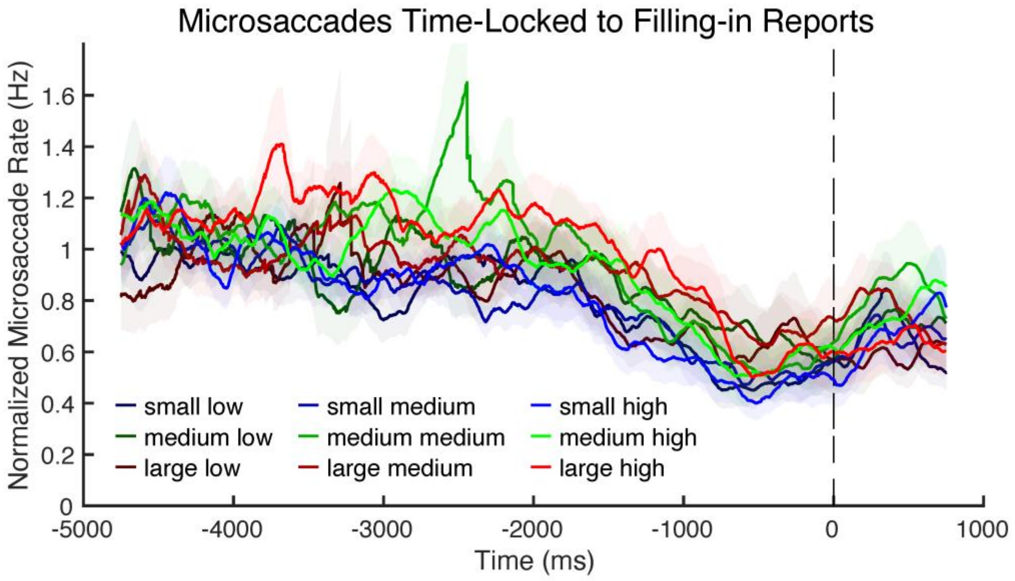


**Fig. S2. Microsaccade rate variability across stimulus conditions.** Data are aligned to the subjective report of perceptual filling-in (time 0). Curves represent different combinations of boundary size (small, medium, large) and color contrast (low, medium, high). Shaded areas indicate standard error of the mean.

**Table S1. Fixed effects from a linear mixed-effect model of filling-in time by stimulus and trial parameters.**

| Parameter | Estimate | 99.17% Percentile CI |
|---|---|---|
| intercept | 0.566 | 0.441, 0.701 |
| contrast | 0.133 | **0.0995, 0.166 *** |
| eccentricity | -0.0930 | **-0.125, -0.0603 *** |
| trial_num | -0.107 | **-0.128, -0.0859 *** |
| contrast : eccentricity | 0.00831 | -0.0319, 0.0526 |
| contrast : trial_num | -0.0203 | -0.0397, 0.000423 |
| eccentricity : trial_num | -0.0116 | -0.0311, 0.00999 |

*: the percentile bootstrap confidence interval (CI), Bonferroni-corrected across fixed effect parameters, does not contain 0 ($\alpha = 0.0083$). The intercept is presented for reference but not evaluated for statistical significance.

**Table S2. Fixed effects from a linear mixed-effect model of filling-in time by stimulus, trial, and eye movement parameters.**

| Parameter | Estimate | 99.29% Percentile CI |
|---|---|---|
| intercept | 0.274 | 0.0994, 0.427 |
| contrast | 0.111 | **0.0810, 0.141** |
| eccentricity | -0.0747 | **-0.0998, -0.0503** |
| trial_num | -0.105 | **-0.122, -0.0865** |
| ms_presence | 0.307 | **0.181, 0.466** |
| num_blinks | 0.274 | **0.210, 0.349** |
| ocular_drift | 0.0293 | -0.0222, 0.0803 |
| contrast : trial_num | -0.0154 | -0.0313, 0.000384 |

This model includes the significant parameters identified in Table S1, plus main effects of eye movements.

*: the percentile bootstrap confidence interval (CI), Bonferroni-corrected across fixed effect parameters, does not contain 0 ($\alpha = 0.0071$). The intercept is presented for reference but not evaluated for statistical significance.
